# Tsetse flies choose birthing sites guided by environmental but not pheromonal cues

**DOI:** 10.1101/2022.11.30.518495

**Authors:** Andrea K. Adden, Lee R. Haines, Álvaro Acosta-Serrano, Lucia L. Prieto-Godino

**Affiliations:** The Francis Crick Institute, London, UK; Liverpool School of Tropical Medicine, Liverpool, UK

## Abstract

Tsetse flies significantly impact public health and economic development in sub-Saharan African countries by transmitting the fatal disease African trypanosomiasis. Unusually, instead of laying eggs, tsetse birth a single larva that immediately burrows into the soil to pupate. Where the female chooses to larviposit is therefore crucial for offspring survival. Previous studies showed that a putative larval pheromone, *n*-pentadecane, attracts gravid female *Glossina morsitans morsitans* to appropriate larviposition sites. However, this attraction could not be reproduced in field experiments. Here, we resolve this disparity by designing naturalistic laboratory experiments that closely mimic the characteristics found in the wild. We show that gravid tsetse were neither attracted to the putative pheromone nor, interestingly, to pupae placed in the soil. In contrast, females appear to choose larviposition sites based on environmental cues. We conclude that it is the substrate, rather than larval pheromones, which drives larviposition site selection under naturalistic conditions.

## Introduction

A wide variety of reproductive strategies have evolved, from a reliance on large numbers of offspring with limited or non-existent maternal care (e.g. mosquitoes, (Shaw *et al*, 2015)), to enormous maternal investment in animals, including our own species, with long pregnancies and small numbers of offspring. This latter strategy has evolved multiple times across phyla (Ostrovsky *et al*, 2016). Tsetse flies (*Glossina* sp.), vectors of the African trypanosome parasites which cause sleeping sickness and animal trypanosomiasis, provide a fascinating example. Instead of laying eggs – like most insects – each tsetse female matures one larva at a time in her uterus. The larva is sustained by feeding on ‘tsetse milk’, which is secreted by modified accessory glands (Denlinger & Ma, 1974). After approximately 10 days *in utero*, when the larva reaches the end of its 3^rd^ instar stage, the female gives birth in a process known as larviposition. The larva then rapidly burrows into the ground on which it was deposited to commence pupation. The female’s choice of larviposition site is thus her final maternal care behaviour, and her choice can markedly affect the offspring’s chance of survival.

One indicator for favourable soil conditions may be the presence of other tsetse larvae and/or pupae, and previous work has suggested that this may be signalled by larval pheromones (Nash *et al*, 1976; Saini *et al*, 1996). Chemical analysis of the larval exudates of two subspecies of *Glossina morsitans* identified the alkanes *n*-pentadecane (for *G. morsitans morsitans*) and *n*-dodecane (for *G. morsitans centralis*) as the major active components for each subspecies, both attracting conspecific gravid females in a two-choice preference assay (Saini *et al*, 1996). Similarly, in *G. brevipalpis* and *G. palpalis*, significant attraction was reported when sand was conditioned with conspecific as well as heterospecific larvae (Renda *et al*, 2016; Gimonneau *et al*, 2021). Although no larviposition pheromone was identified in these studies, the authors suggested that pheromones may be similar across these two species.

Given that tsetse have a significant impact on public health and economic development in sub-Saharan African countries (Kuzoe & Schofield, 2004; Alsan, 2015), a larviposition pheromone that reliably attracts gravid females would open promising new avenues in vector control. However, when a recent study tested the effectiveness of *n*-pentadecane as an attractant of gravid female tsetse in the field, it found no effects at any concentration (Hargrove *et al*, 2021; see also Rowcliffe & Finlayson, 1981). These contradictory results raise the question of whether findings from behavioural assays, performed under artificial laboratory conditions, are applicable to the complex natural ecology of tsetse in the natural environment. We aimed at closing this gap, and revisiting the evidence for the existence of a larviposition pheromone in *G. m. morsitans*, by conducting carefully controlled laboratory experiments that nevertheless presented the flies with conditions that were as naturalistic as possible – that is to say they matched as closely as possible the qualities of larviposition sites that a fly might encounter in the field.

Our results demonstrate that tsetse display clear preference for certain environmental cues, such as substrate consistency and shade, which is in agreement with studies that surveyed pupation sites in the field (Phelps & Burrows, 1969). However, we found that under naturalistic settings, neither the putative *G. m. morsitans* pheromone *n*-pentadecane nor the aggregation of pupae in the substrate had any effect on the larviposition choices of these flies. We conclude that it is unlikely that larviposition in *G. morsitans* is guided by larval pheromones in natural ecological settings, and that other environmental cues are the major drivers of female larviposition choices.

## Results

### Minimising stress in larviposition choice assays

Previous studies employed standarised laboratory methods that have the potential to induce stress on tsetse flies, yet they did not asses whether flies were stressed and if this could impact the results. In addition to the handling of the gravid females, the two most important stressors introduced in these assays were the colour of the behavioural arenas and the larviposition substrate. Two out of three studies (Saini *et al*, 1996; Renda *et al*, 2016) used white enclosures, and all studies used plain white sand as a larviposition substrate (Saini *et al*, 1996; Renda *et al*, 2016; Gimonneau *et al*, 2021). Bright spaces are, however, usually associated with approaching host animals, whereas females in late stages of pregnancy rarely feed, and are more likely to rest in shaded areas (Pilson & Pilson, 1967). Furthermore, field work has shown that tsetse tend to larviposit under fallen logs, in tree holes and in underground burrows, and prefer leaf litter to exposed sand (Muzari & Hargrove, 2005; Rowcliffe & Finlayson, 1981; Phelps & Burrows, 1969).

To address these discrepancies, our behavioural paradigm was designed to be as naturalistic and stress-free for the flies as possible, while also being a carefully controlled insectary experiment. Non-forced two-choice experiments were conducted in a large flight cage with brown-stained netting, and we used natural colour sand (Fig. S1A, B).

We evaluated female stress by (1) counting the number of abortions induced by handling, (2) calculating the proportion of females that larviposited within 48 hours (larviposition rate), and (3) measuring the weight of the pupae as an indicator of sustained female investment (Fig. 1). We tested a total of 1,639 female *G. m. morsitans*, which were selected for experiments if the polypneustic lobes of the larva were visible through the female’s cuticle (Fig. 1A). The females produced 1,423 larvae within 48h, at an overall larviposition rate of 86.8%, which was highly consistent across experimental conditions (Fig. 1B). The overall abortion rate was a negligible 0.43%, and the mortality rate among females was <2%.

**Figure 1:**
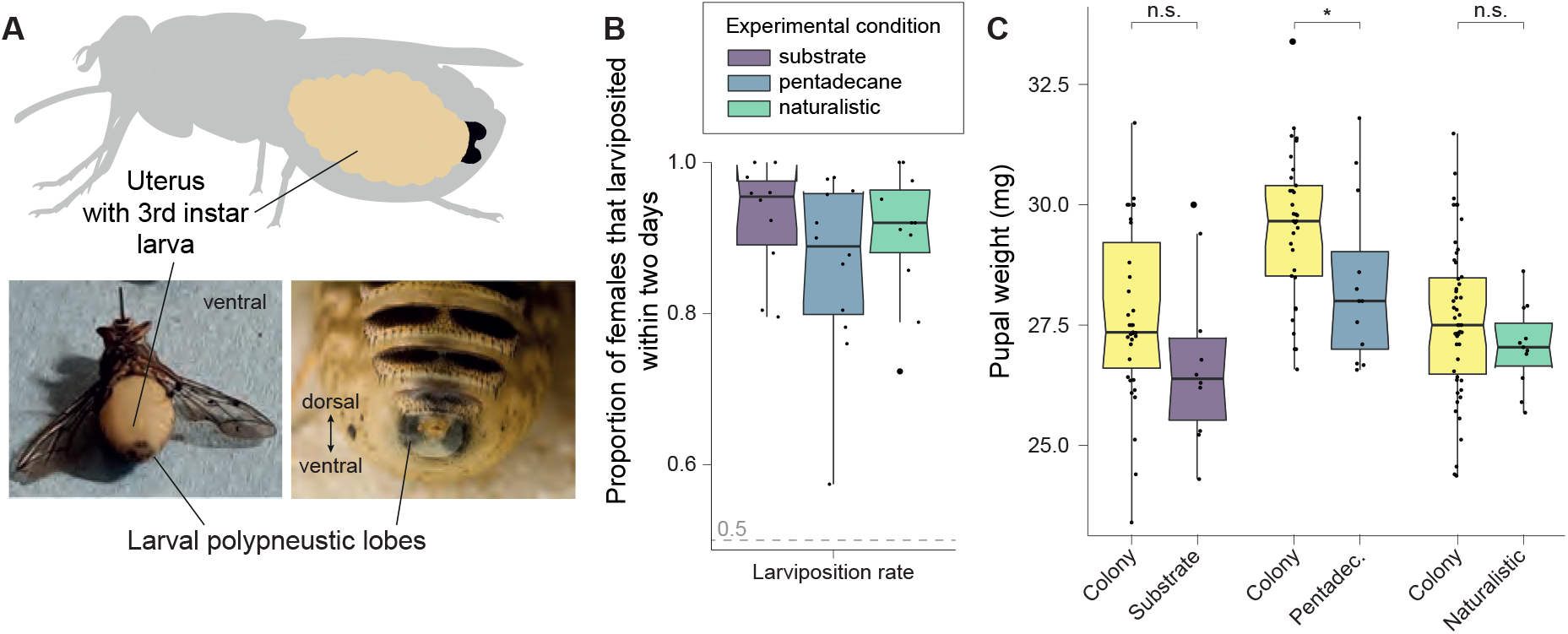
Female and larval health. (A) Females were selected for experiments if the polypneustic lobes of the larva were clearly visible through the female’s cuticle. (B) Proportion of females that larviposited within two days (larviposition rate). Median larviposition rates for the three experimental conditions were 95.46% (‘substrate’), 88.88% (‘pentadecane’) and 92.00% (‘naturalistic’). The larviposition rate was highly consistent across experimental conditions (n.s. Kolmogorov-Smirnov test). (C) Pupal weight measured for all experimental conditions and compared to Pupal weights measured in the colony at the time of the experiment. Median Pupal weights were 26.39 mg (subs.), 28.0 mg (pent.) and 27.01 mg (nat). The distributions of Pupal weights obtained from pentadecane experiments are significantly smaller from colony weights (p = 0.0233, Kolmogorov-Smirnov test). However, all experimental weight distributions fall within the expected weight distribution of the colony.

As a measure of larval health, we measured the weight of all deposited pupae. For all experiments, pupal weight distributions fell within the measured distribution of the colony at the time of the experiment (Fig. 1C), although experimental pupae were slightly smaller than control pupae. However, colony records show that first-time mothers produce overall smaller larvae than older mothers, and females chosen for our experiments often included first-time mothers (Lord *et al*, 2021). It is therefore not unexpected that median experimental weights were lower. We conclude that stress was not a significant factor in our experiments.

### Tsetse prefer to larviposit on leaf litter over sand

Field experiments have reported that tsetse prefer to larviposit on leaf litter over sand (Muzari & Hargrove, 2005), yet all prior laboratory larviposition assays were done using sand as a substrate (Saini *et al*, 1996; Renda *et al*, 2016; Gimonneau *et al*, 2021). To confirm which substrate is preferred by tsetse, we tested females in a non-forced two-choice larviposition assay, in which flies could choose to larviposit in trays filled either with sterilised leaf litter or sand, or on the floor (Fig 2C). We found that when corrected for surface area, flies preferred to larviposit on the trays over the floor, and this held true across all experiments (Fig. 2B). Because we were interested in the choice between the two substrates, we only used larvae deposited on either tray to calculate preference indexes (PI). We found a clear preference for leaf litter, with a median PI of 0.82 (Fig. 2D) and significantly higher larval counts in the leaf litter tray as compared to the sand tray (p < 0.001, Wilcoxon rank sum test; Fig. 2E). Across trials, 90.46% of larvae were deposited on leaf litter. We therefore conclude that gravid tsetse females prefer to larviposit on leaf litter, as suggested by previous reports from the field. This highlights that previous larviposition experiments, carried out in the laboratory, were performed with suboptimal substrates. Given these results, we chose to use leaf litter rather than sand as larviposition substrate in the following experiments.

**Figure 2:**
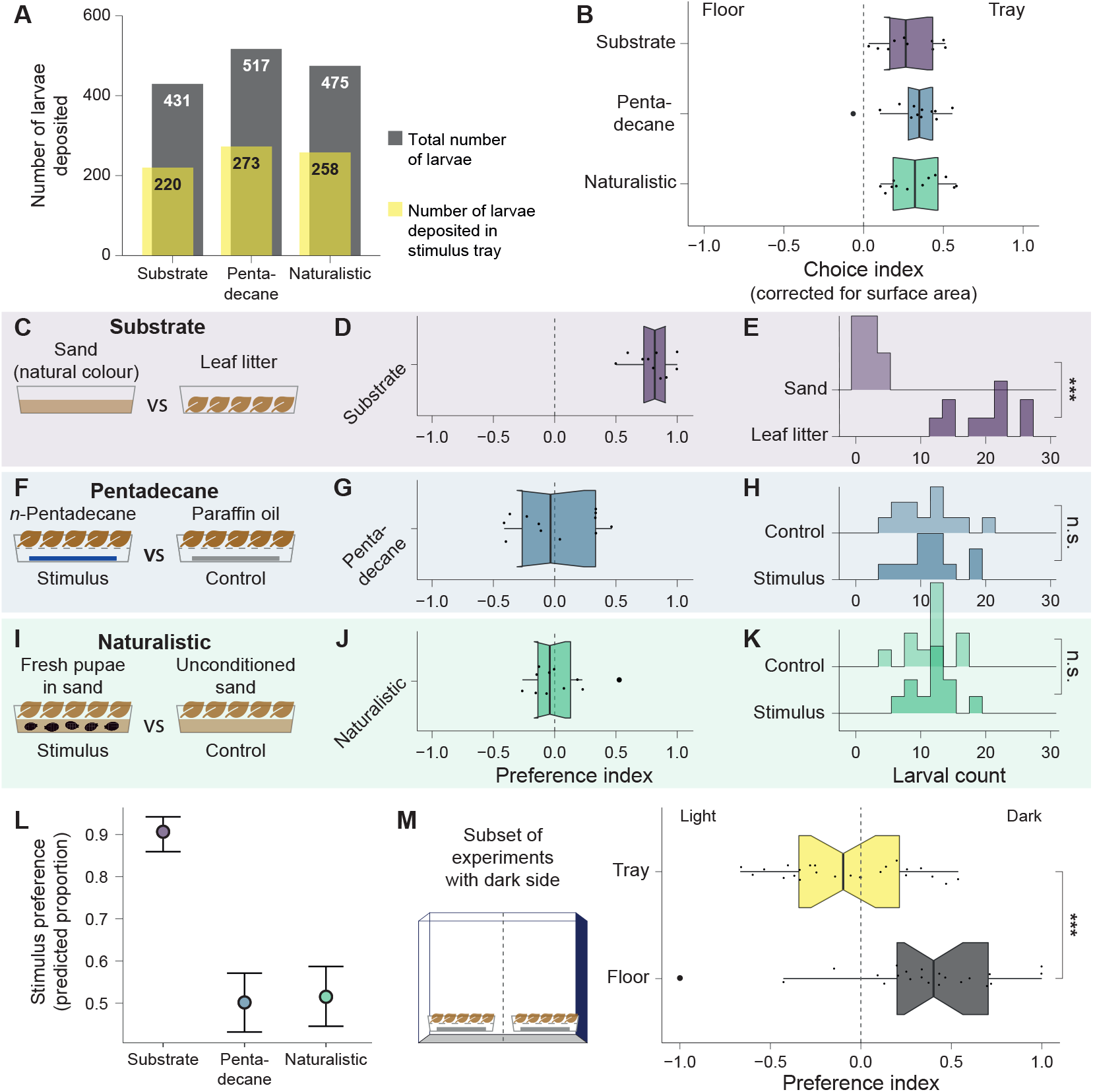
Female larviposition preferences. (A) Absolute numbers of larvae deposited during experiments. Grey: Total number of larvae deposited across cages per experiment. Yellow: Number of larvae deposited in stimulus trays, rather than on the floor. Larvae deposited on the floor were not included in the analysis. (B) Choice index for larvipositing on trays vs. on the floor. The choice index was corrected for surface area. Across experiments females prefer to larviposit on trays. (C) In the ‘substrate’ condition, females were offered the choice between natural colour sand and leaf litter as larviposition substrate. (D) The preference index for the ‘substrate’ condition shows a strong preference for leaf litter (median PI = 0.817). (E) A ridge histogram of larval count for each stimulus in the ‘substrate’ condition shows a significantly higher larval count in leaf litter (p < 0.001, Wilcoxon rank sum test). (F) In the ‘pentadecane’ condition, females chose between the proposed larval pheromone *n*-pentadecane, and the solvent paraffin oil as a control stimulus. (G) Preference index for the ‘pentadecane’ condition shows no preference for *n*-pentadecane. (H) The distributions of larval count in the stimulus and control trays strongly overlap. (I) In the ‘naturalistic’ condition, females chose between sand conditioned with 20 1-day old pupae, and unconditioned sand. (J) Preference index for the ‘naturalistic’ condition showns no preference for conditioned sand. (K) The distributions of larval count in conditioned vs. unconditioned sand strongly overlap. (L) Conditional effects predicted by a Bayesian model of female choice. The model supports a preference only in the ‘substrate’ condition (estimated median = 0.906, standard error = 0.02) but not in the ‘pentadecane’ or ‘naturalistic’ conditions (estimated medians 0.502 and 0.5125, respectively). (M) In a subset of experiments, the cage had one dark wall. The preference index for cage side shows that females depositing in trays do not have a preference for the darker side of the cage (no difference from normal distribution, mean=0, sd=1, Kolmogorov-Smirnov test), but females depositing on the floor prefer the darker half of the floor (median PI = 0.402; difference from normal distribution, mean=0, sd=1: p << 0.001, Kolmogorov-Smirnov test).

### *n*-pentadecane does not act as a larviposition pheromone under naturalistic laboratory settings

We next tested whether the described larval pheromone *n*-pentadecane attracts females for larviposition. We presented a dilution of *n*-pentadecane on filter paper that was the equivalent of volatile released by 80 pupating larvae over 2 hours – a dose that was reported to attract approximately 75% of females (Saini *et al*, 1996). The *n*-pentadecane was presented on filter paper under a suspended layer of leaf litter, such that the females could not touch the *n*-pentadecane itself (Fig. 2F). As a control stimulus, the solvent paraffin oil was also presented on filter paper under leaf litter. Under these conditions, we found no preference (median PI = -0.034, Fig. 2G), with 50.18% of larvae deposited in the stimulus tray. Furthermore, the distributions of larval counts in each stimulus tray strongly overlap (Fig. 2H). This suggests that *n*-pentadecane alone is not sufficient to attract females to a larviposition site.

### No evidence for the presence of a larviposition pheromone in tsetse

We reasoned that although *n*-pentadecane did not act as a larviposition pheromone, other compounds emitted by pupating larvae might contribute to an attractive effect. To test for a putative larviposition pheromone, we devised a naturalistic experiment in which 20 one-day old pupae were presented in sand (where they had pupated) under a layer of leaf litter, again so the flies were unable to make contact with the sand (Fig. 2I). The control stimulus was sand that did not contain any pupae. Under these conditions, females again showed no preference (PI = -0.04, Fig. 2J), with 50.78% of pupae depositing in the stimulus tray. As in the ‘pentadecane’ condition, the larval count distributions strongly overlap (Fig. 2K). We therefore conclude that under naturalistic conditions, larval and pupal volatiles are not sufficient to attract females to a larviposition site. These results are supported by a categorical model of experimental conditions using Bayesian inference, which supports a preference only in the ‘substrate’ condition but not in the ‘pentadecane’ or ‘naturalistic’ conditions (Fig. 2L).

### Shade is preferred but is secondary to substrate quality

Another cue that has been proposed to be important for larviposition preference in the field is the presence of shade (Muzari & Hargrove, 2005). Although we did not test this directly, one set of experiments was carried out in cages where one cage wall was darker than the others (see Methods). When we analysed these results, we found that the number of larvae deposited on the trays filled with leaf litter did not depend on whether the trays were oriented to the dark or light side of the cage (Fig. 2M). However, for larvae deposited on the floor, there was a clear preference for the dark side of the cage (Fig. 2M). These results indicate that while substrate quality is a key larviposition cue, a darker environment is preferred in the absence of other cues, something that has also been suggested previously (Rowcliffe & Finlayson, 1981).

## Discussion

An outstanding question in tsetse ecology was whether these viviparous flies use pheromones to guide their larviposition choices. Three laboratory studies argued this to be the case, and one study identified the putative pheromone compound for *G. m. morsitans, n*-pentadecane (Saini *et al*, 1996; Renda *et al*, 2016; Gimonneau *et al*, 2021). However, a recent study in the field was unable to reproduce these results, arguing that such putative compounds do not act as pheromones in tsetse natural ecology (Hargrove *et al*, 2021). Here we bridge the gap between larviposition experiments in the laboratory and in the field. We have shown that it is possible to undertake naturalistic larviposition experiments under tightly controlled laboratory conditions, allowing us to reproduce results from the field. Our results suggest that the prior discrepancies between laboratory and field studies may be due to artificial conditions imposed on gravid females in laboratory settings. Importantly, our work paves the way for future behavioural experiments in the laboratory under more naturalistic conditions.

We show that under naturalistic conditions in the laboratory, pheromones do not to play a role in female larviposition choice, and that environmental cues are the main drivers of this choice. Among environmental cues, larviposition substrate seems to be the most important one, with tsetse clearly preferring to larviposit on leaf litter over sand. Furthermore, we show that shaded environments are also preferred, which is consistent with prior field data (Hargrove *et al*, 2021; Muzari & Hargrove, 2005). Leaf litter has a well-described function of shielding the soil surface from rapid changes in temperature and humidity, thus helping to create a stable environment for soil-living organisms (Sayer, 2006). Rates of pupal development in tsetse are strongly temperature-dependent (Phelps, 1973; Hargrove & Vale, 2020) and gravid females are known to seek out shaded and sheltered areas such as underground warthog burrows to deposit their larvae (Muzari & Hargrove, 2005) where temperature fluctuations are minimized.. The presence of leaf litter alone is therefore likely to signal appropriate larviposition sites, both in terms of maternal care by selecting optimal conditions for larval development, and in providing shade and shelter for the female and her larva during parturition.

While we confirm that gravid females strongly prefer leaf litter, previous laboratory studies traditionally provided sand as a larviposition substrate (Saini *et al*, 1996; Gimonneau *et al*, 2021; Renda *et al*, 2016). The sand used in these studies was white, in contrast to the natural-coloured sand we used in the present study. It is well described that tsetse are not attracted to white surfaces (Brady & Shereni, 1988; Lindh *et al*, 2012). Therefore, the brightness of the white sand may have added to the artificial nature of the experiments. In the field, flies tend to rest in the shade, e.g. under tree branches, throughout the hottest part of the day (Pilson & Pilson, 1967). We conclude that factors such as shade can strongly influence the flies in the absence of other cues, as previously reported (Rowcliffe & Finlayson, 1981). It is clear that in future studies, the light/shade distribution in the cages needs to be taken into account and, if possible, controlled for.

Our finding that pheromones do not guide larviposition decisions under naturalistic conditions can be explained by a number of factors in the ecological niche that tsetse inhabit. Although a pheromone could mark a site at which other larvae have successfully pupated, thus overcoming environmental obstacles such as desiccation and overheating (Hargrove & Vale, 2020), *n*-pentadecane is only released during early pupariation (Nash *et al*, 1976). As pupal development takes approximately 3-4 weeks in *G. morsitans, n*-pentadecane would carry no information about the subsequent survival of the pupae. Furthermore, tsetse pupae are at constant risk of predation by ants and beetles (Fiske, 1920; Adabie, 1987; Rogers & Randolph, 1990), and a volatile pheromone strong enough to attract females may also attract predators. While there could be strength in numbers when it comes to creating a microenvironment optimal for the pupae to survive and develop in, it has also been shown that this is only the case up to a certain pupal density, beyond which aggregation becomes detrimental to survival due to predation (Rogers & Randolph, 1984). Thus, the advantages of a larval pheromone remain unclear. Additionally, in the field, leaf litter releases a substantial number of volatiles, potentially including *n*-pentadecane – a known volatile of many plant species. We therefore favour the proposal of (Hargrove *et al*, 2021) that in natural settings, plant pentadecane will likely mask larval volatiles. As a consequence, larval *n*-pentadecane emissions may be of practical use only under very specific circumstances, e.g. when larviposition sites are limited and bare soil or sand are the only options. This is in agreement with multiple fieldwork studies that have established that in the wild tsetse select larviposition sites based on season, temperature, and soil conditions (Fiske, 1920; Muzari & Hargrove, 2005; Rowcliffe & Finlayson, 1981; Adabie, 1987).

To conclude, this study demonstrates the importance of taking an animal’s natural environment into account when designing laboratory experiments with colony-reared insects. In the case of the tsetse, new attractants found in the laboratory understandably create much interest as potential new ways of vector control. While carefully controlled laboratory studies can generate exciting new insights into tsetse biology, these insights may not be relevant when taking the flies’ ecology and natural environment into account, and attractants identified in the lab may indeed be overridden by other environmental cues. We therefore recommend that future insectary-based studies match the natural environment as far as possible.

## Materials and methods

### Animals

*Glossina morsitans morsitans* Westwood were bred at the Liverpool School of Tropical Medicine, UK. Tsetse rooms were kept at a temperature of 26°C-28°C and 60-80% humidity at all times, with a 12h/12h light/dark cycle. The health of all female flies involved in experiments was monitored by recording abortion rates and the weights of offspring (pupae). Females were selected for participation in an experiment when the polypneustic lobes of the larva *in utero* were visible through the female’s posterior cuticle (Fig. 1D), indicating that parturition would occur within the following 24-48 hours. For selection purposes, females were kept in a cold room at 2-6°C for no longer than 10 minutes and were allowed to warm up in the colony room before being transferred to the experimental cage. Each female was used once per experimental condition. We used females of varying ages (between 16 and 60 days old, i.e. between their first and fifth gravidity cycle) to reflect the natural situation as closely as possible. After participating in experiments, females readily fed and continued to produce larvae until they were terminated at 10 weeks old.

### Behavioural experiments

All experiments were done in 60×60×90 cm pop-up insect cages (Watkins & Doncaster, Leominster, Herefordshire, UK). Cages were initially washed with bleach, and subsequently stained brown using black tea (Unilever UK PG Tips, London, UK). One wall (60×60 cm) of the cage was dark blue, which we used as the floor in the ‘substrate’ and ‘naturalistic’ conditions. In the ‘pentadecane’ condition, the cage was placed horizontally (60×90 cm floor area).

Leaf litter was collected at Hampstead Heath (London, UK), sterilised in an autoclave for two hours at 121°C and dried at 100°C for one hour. Between experiments, leaf litter was re-sterilised using a drying oven at 121°C for two hours. Natural colour silica sand (Trustleaf, March, UK) was washed in tap water and then sterilised in an oven at 121°C for four hours. Between experiments, sand was sterilised again at 121°C for four hours. Natural colour sand was chosen to match the leaf litter colours better than white sand to avoid presenting a visually distinct stimulus. Stimuli were presented in 28×17.5×17 cm large food-grade plastic trays (Solent Plastics, Romsey, UK), which each covered approximately 30.7% of the available floor space (Fig. S1).

Experiments were run over a two-day period, during which time approximately 80% of females larviposited. On the morning of the second day, stimulus and control trays were exchanged for fresh trays. Stimulus and control trays were placed on the right or left side randomly on day 1, and in reverse order on day 2. Note that we took care to finish these preparations by 13:00h, so as not to interfere with the flies’ period of peak larviposition, which occurred between 14:00h and 18:00h. Flies were then left unattended for the remainder of the day to keep human presence in the room to a minimum, as savannah tsetse species, particularly females, can be repelled by human odour (Vale, 1974; Torr *et al*, 2012).

Pupae deposited in trays were counted every day. To be able to count pupae deposited on the floor, double-sided gel tape (Nano-Grab, Shurtape Technologies, Avon, OH, US) was used as a separation between the right and left half of the cage floor (Fig. S1). Pupae were often found stuck to the side of the tape, indicating that the larvae could not cross this barrier. We therefore assumed that pupae found on each side were deposited there, and counted them accordingly.

After two days, experiments were terminated and all flies were removed from the cages using a fly aspirator (Katcha Bug Buster, Select IP Ltd, Chesterfield, UK).

### Testing the substrate: sand vs. leaf litter

To test whether the flies prefer sand or leaf litter as a substrate for larviposition, we first presented 800 mL sterilised sand and 800 mL sterilised leaf litter to gravid female *G. m. morsitans*. Pupae were counted daily. The experiment was repeated 10 times, with 39-51 females added per cage, amounting to a total of 465 females.

### Testing the pheromone: n-pentadecane vs. paraffin oil

According to Saini et al. (1996), *n*-pentadecane was the primary active compound of the larviposition pheromone of *G. m. morsitans*. We aimed at replicating this finding in the lab, but using leaf litter as a larviposition substrate. To this end, we used a 10^−2^ dilution of *n*-pentadecane (Sigma-Aldrich, Steinheim, Germany; Lot# BCCC7129) in paraffin oil, with a resulting concentration of 7.69 μg/μL. Using 25 μL of this dilution delivered a dose of *n*-pentadecane equivalent to approx. 192 μg – equivalent, approximately, to the amount released by 80 pupating larvae over two hours (Saini et al. 1996). According to Saini and colleagues, this dose should be sufficient to show robust attraction of approximately 75% of females, while not being too high to act as a repellent. As a control stimulus, we used 25 μL of paraffin oil.

Both stimuli were pipetted onto discs of filter paper (Whatman No.1) and placed in the bottom of a stimulus tray. The tray was then covered with a cardboard-framed mesh that allowed burrowing larvae to fall into the tray below, but prevented flies from entering the tray. Approximately 1 L of leaf litter was then placed on top of the mesh. Stimuli were placed randomly on the left or right side of the cage, and exchanged for fresh stimuli on day 2 as detailed above. Pupae were collected and counted as described above. The experiment was repeated 12 times, amounting to a total of 601 females tested (between 45-55 females per cage).

### Naturalistic experiments

To confirm our findings and make them comparable to results obtained from field studies, we conducted an additional experiment aimed at being as close to a natural situation as possible. First, we reasoned that the earliest-depositing female flies on any given day would not encounter freshly pupated larvae in the ground – rather, they would find an area in which larvae deposited the previous day are respirating. Second, the ground would likely be covered in leaf litter or other organic debris. Finally, any additional larva deposited on the same day would add to the complex bouquet of larval and pupal odours that may play a role in attracting females to the site. We therefore designed the following experiment.

Between 14:00h and 18:00h (peak larviposition time), 20 freshly deposited larvae were collected and placed in a tray containing 800 mL sterilised sand. Most larvae quickly burrowed into the sand, while a small minority pupated on top of the sand. A control tray of larvae-free sand was prepared at the same time. Both trays were covered and left in the fly colony room overnight. The following day (day 2), mesh frames were placed onto both trays, which were subsequently covered in 800 mL sterilised leaf litter. The leaf litter was separated from the sand in this way to prevent flies from touching the sand. Trays were placed into an experimental cage containing 41-52 heavily pregnant females. Also on this day, new stimulus and control trays were prepared by adding larvae to sand as described above.

On day 3, stimulus and control trays were exchanged for freshly made trays and the position of the stimulus and control trays was reversed. All pupae were removed from the trays and from the floor of the cage and counted. On the final day (day 4), trays were removed and all pupae counted. All flies were removed from the cages using a fly aspirator. In total, 573 females were tested in this experimental condition.

### Shade experiment

To gain insight into other factors that may contribute to the females’ larviposition choice, we took advantage of the fact that we had conducted the ‘pentadecane’ experiment placing the cage horizontally, with one side of the cage flanked by a dark blue cage wall. It is well documented that tsetse are attracted to blue surfaces, and we aimed to test whether this experimental arrangement impacted larviposition choices. Stimulus and control trays were placed on alternating sides of the cage as described above, but for analysis purposes, pupae were grouped according to the half of the cage in which they were recovered – that is, the ‘dark’ side or the ‘light’ side. The analysis was performed for pupae found in the trays and on the floor.

### Statistical analysis

Pupae deposited in each cage cannot be assumed to be independent, as the presence of a pupa in a stimulus tray may influence the larviposition decision of other females in the cage. We therefore calculated a preference index per cage, given as

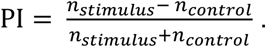

Thus, PI = 1 indicates that females larviposited exclusively in the stimulus tray, while PI = -1 indicates that all females preferred the control. PI = 0 indicates no preference.

The choice index refers to the preference of females to larviposit on stimulus trays or on the floor. However, as the total available surface area on the floor was larger than on the trays, it was calculated based on the density of pupae found on the floor vs. the trays, in pupae/cm^2^. The preference index for the light or shaded half of the cage was calculated in an analogous way to the preference index described above, contrasting the number of pupae found on trays and on the floor in the right (shaded) vs. left (light) half of the cage.

To compare distributions of larvae across cages within one experimental condition, we used the non-parametric Wilcoxon rank sum test. Across experiments, we compared distributions using the Kolmogorov-Smirnov test. Analysis of variance (Anova) was used to test the effects of other factors that may have influenced the results.

For further statistical analysis, we built a Bayesian model using the R-package *brms* (Bürkner, 2017). Samples were drawn from a Bernoulli-type distribution, using default priors in addition to weakly informative priors for slope (normal distribution with mean 0, standard deviation 5), with leave-one-out cross-validation (LOO-CV) as an added criterion. We modelled female choice, binarised to 1 (female chose stimulus/leaf litter) and 0 (female chose control/sand), for the three experimental conditions, using only cage as a group-level effect. The model was run across four chains with 1000 warm-up and 4000 post-warmup draws each. The chains converged according to Rhat values, effective sample size values, and visual representations of chain traces. For further information, see supplementary file S3.

All statistical analysis was done in RStudio using R 4.2.1. Data was plotted using *ggplot2* (Wickham, 2016) and *viridis* (Garnier *et al*, 2021). Figures were prepared in Adobe Illustrator 24.0.1.

#### Statement of authorship

AKA, LRH and LPG designed the study. ÁAS and LRH provided the animals and facilities. AKA conducted the experiments, analysed the data, prepared the figures and wrote the first draft of the manuscript. All authors contributed to the final version of the manuscript.

## Acknowledgements

The authors would like to thank Mr. Zach Stavrou-Dowd and Dr. Clair Rose for helping maintain the tsetse colony, and John Kirwan for assistance with the Bayesian modelling. AKA was funded by an EMBO long-term fellowship; LRH was funded by a Wellcome Trust Institutional Strategic Support Fund (grant 204806/Z/16/Z, internal award reference DCF19031921LH); ÁAS was partially supported by UK Medical Research Council (MRC) Confidence in Concept award 2017-18 MC_PC_17167, and Biotechnology and Biological Sciences Research Council (BBSRC) Anti-VeC award AV/PP0021/1; and LLPG’s laboratory is supported by a European Research Council (ERC) Starting Investigator Grant (802531), an Allen Distinguished Investigator Award, a Human Frontiers Science Grant (RGY0052/2022) and The Francis Crick Institute, which receives its core funding from Cancer Research UK (CC001594), the UK Medical Research Council (CC001594) and the Wellcome Trust (CC001594).

## References

Adabie DA (1987) Pupal ecology and role of predators and parasitoids in natural population regulation of Glossina pallidipes Austen (Diptera: Glossinidae) at Nguruman, Kenya.

Alsan M (2015) The Effect of the TseTse Fly on African Development. Am Econ Rev 105: 382–410

Brady J & Shereni W (1988) Landing responses of the tsetse fly Glossina morsitans morsitans Westwood and the stable fly Stomoxys calcitrans (L.) (Diptera: Glossinidae & Muscidae) to black-and-white patterns: A laboratory study. Bull Entomol Res 78: 301– 311

Bürkner P-C (2017) brms: An R Package for Bayesian Multilevel Models Using Stan. J Stat Softw 80: 1–28

Denlinger DL & Ma W-C (1974) Dynamics of the pregnancy cycle in the tsetse Glossina morsitans. J Insect Physiol 20

Fiske WF (1920) Investigations into the bionomics of glossina palpalis. Bull Entomol Res 10: 347–463

Garnier S, Ross N, Rudis R, Camargo AP, Sciaini M & Scherer C (2021) viridis - Colorblind-Friendly Color Maps for R.

Gimonneau G, Romaric O, Salour E, Rayaisse J-B, Buatois B, Solano P, Dormont L, Roux O & Bouyer J (2021) Larviposition site selection mediated by volatile semiochemicals in Glossina palpalis gambiensis. Ecol Entomol 46: 301–309

Hargrove JW, Van Sickle J & Saini RK (2021) Field Testing of a Putative Larviposition Pheromone for the Tsetse Fly Glossina m. morsitans Westwood. African Entomol 29: 635–648

Hargrove JW & Vale GA (2020) Models for the rates of pupal development, fat consumption and mortality in tsetse (Glossina spp). Bull Entomol Res 110: 44–56

Kuzoe FAS & Schofield CJ (2004) Strategic Review of Traps and Targets for Tsetse and African Trypanosomiasis Control. World Heal Rep: 1–58

Lindh JM, Goswami P, Blackburn RS, Arnold SEJ, Vale GA, Lehane MJ & Torr SJ (2012) Optimizing the colour and fabric of targets for the control of the tsetse fly Glossina fuscipes fuscipes. PLoS Negl Trop Dis 6

Lord JS, Leyland R, Haines LR, Barreaux AMG, Bonsall MB, Torr SJ & English S (2021) Effects of maternal age and stress on offspring quality in a viviparous fly. Ecol Lett: 1– 12

Muzari MO & Hargrove JW (2005) Artificial larviposition sites for field collections of the puparia of tsetse flies Glossina pallidipes and G. m. morsitans (Diptera: Glossinidae). Bull Entomol Res 95: 221–229

Nash TAM, Trewern MA & Moloo SK (1976) Observations on the free larval stage of Glossina morsitans morsitans Westw. (Diptera, Glossinidae): The possibility of a larval pheromone. Bull Entomol Res 66: 17–24

Ostrovsky AN, Lidgard S, Gordon DP, Schwaha T, Genikhovich G & Ereskovsky A V. (2016) Matrotrophy and placentation in invertebrates: a new paradigm. Biol Rev Camb Philos Soc 91: 673–711

Phelps RJ (1973) The effect of temperature on fat consumption during the puparial stages of Glossina morsitans morsitans Westw. (Dipt., Glossinidae) under laboratory conditions, and its implication in the field. Bull Entomol Res 62: 423–438

Phelps RJ & Burrows PM (1969) Prediction of the pupal duration of Glossina morsitans orientalis Vanderplank under field conditions. J Appl Ecol 6: 323–337

Pilson RD & Pilson BM (1967) Behaviour studies of Glossina morsitans Westw. in the field. Bull Entomol Res 57: 227–257

Renda S, De Beer CJ, Venter GJ & Thekisoe OMM (2016) Evaluation of larviposition site selection of Glossina brevipalpis. Vet Parasitol 215: 92–95

Rogers DJ & Randolph SE (1984) A review of density-dependent processes in tsetse populations. Int J Trop Insect Sci 5: 397–402

Rogers DJ & Randolph SE (1990) Estimation of rates of predation on tsetse. Med Vet Entomol 4: 195–204

Rowcliffe C & Finlayson LH (1981) Factors influencing the selection of larviposition sites in the laboratory by Glossina morsitans morsitans Westwood (Diptera: Glossinidae). Bull Entomol Res 71: 81–96

Saini RK, Hassanali A, Andoke J, Ahuya P & Ouma WP (1996) Identification of major components of larviposition pheromone from larvae of tsetse flies Glossina morsitans morsitans Westwood and Glossina morsitans centralis Machado. J Chem Ecol 22: 1211–1220

Sayer EJ (2006) Using experimental manipulation to assess the roles of leaf litter in the functioning of forest ecosystems. Biol Rev Camb Philos Soc 81: 1–31

Shaw WR, Attardo GM, Aksoy S & Catteruccia F (2015) A comparative analysis of reproductive biology of insect vectors of human disease. Curr Opin Insect Sci 10: 142– 148

Torr SJ, Chamisa A, Mangwiro TNC & Vale GA (2012) Where, when and why do tsetse contact humans? Answers from studies in a national park of Zimbabwe. PLoS Negl Trop Dis 6

Vale GA (1974) The responses of tsetse flies (Diptera, Glossinidae) to mobile and stationary baits. Bull Entomol Res 64: 545–588

Wickham H (2016) ggplot2: Elegant Graphics for Data Analysis Springer-Verlag New York

