## Supplementary information for "Tsetse flies choose birthing sites guided by environmental but not pheromonal cues"

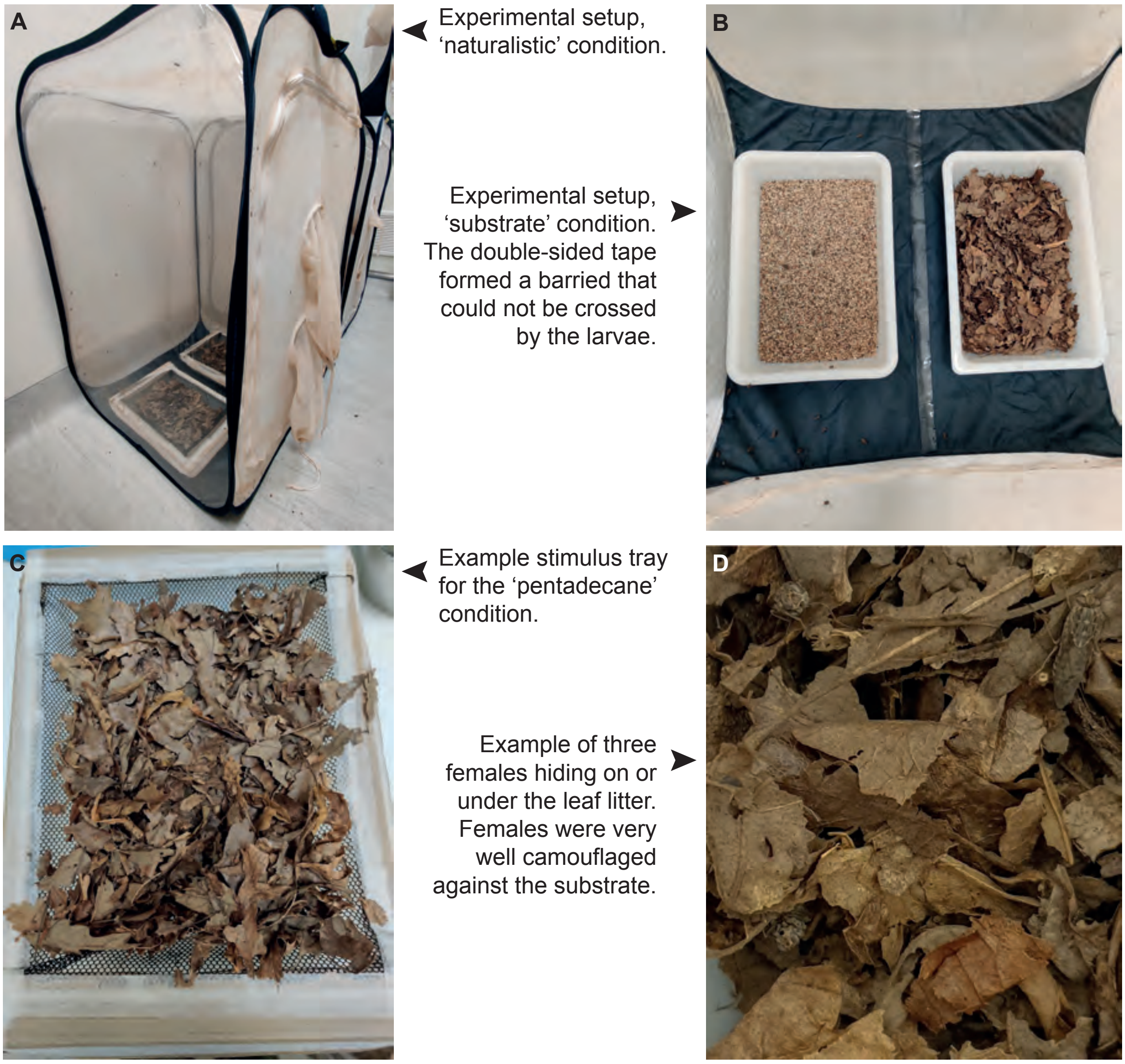


**Figure S1: Photos of the experimental setup.** (A) Experimental setup for the 'naturalistic' condition. (B) Experimental setup for the 'substrate' condition, showcasing the double-sided tape that formed a barrier between the two halves of the experimental cage. (C) Example stimulus tray for the 'pentadecane' condition, where filter paper containing the stimulus or control odour was hidden under a loose layer of leaf litter. (D) Example of three females hiding in leaf litter.

### S2: brms analysis summary

2022-11-08

#### Bayesian model for female larviposition choice analysis.

Import libraries:

```
## Loading required package: Rcpp

## Loading 'brms' package (version 2.17.0). Useful instructions
## can be found by typing help('brms'). A more detailed introduction
## to the package is available through vignette('brms_overview').

##
## Attaching package: 'brms'

## The following object is masked from 'package:stats':
##
##     ar

##
## Attaching package: 'dplyr'

## The following objects are masked from 'package:stats':
##
##     filter, lag

## The following objects are masked from 'package:base':
##
##     intersect, setdiff, setequal, union
```

##### Load the data

```
d_sub <- read.csv('/Users/addena/Documents/R/LSTM/Sand_vs_leaflitter_August.csv',
                  sep=",") # substrate experiment
d_nat <- read.csv('/Users/addena/Documents/R/LSTM/naturalistic_exp.csv',
                  sep=",") # naturalistic experiment
d_pent <- read.csv('/Users/addena/Documents/R/LSTM/C15_vs_PO_data.csv',
                  sep=",") # pentadecane experiment
```

##### Condense the data into one data frame

We remove cage 7 from the 'naturalistic' condition as it was terminated prematurely.

```r
d_sub <- aggregate(cbind(sand, leaf) ~ cage, data = d_sub, FUN = sum, na.rm = T)
d_sub$experiment <- c(rep('3_substrate', 10))
d_sub <- rename(d_sub, stim = leaf)
d_sub <- rename(d_sub, con = sand)

d_nat <- aggregate(cbind(stim, con) ~ cage, data = d_nat, FUN = sum, na.rm = T)
d_nat$experiment <- c(rep('1_naturalistic', 12))
d_nat <- d_nat[-7,] # remove cage 7 (terminated prematurely)

d_pent <- aggregate(cbind(stim, con) ~ cage, data = d_pent, FUN = sum, na.rm = T)
d_pent$experiment <- c(rep('2_pentadecane', 12))

data <- rbind(d_sub, d_nat, d_pent)
data
```

```
##     cage con stim     experiment
## 1      1   2   15    3_substrate
## 2      2   2   22    3_substrate
## 3      3   5   20    3_substrate
## 4      4   0   23    3_substrate
## 5      5   3   18    3_substrate
## 6      6   1   14    3_substrate
## 7      7   0   26    3_substrate
## 8      8   1   22    3_substrate
## 9      9   4   12    3_substrate
## 10    10   3   27    3_substrate
## 13     1  12   19 1_naturalistic
## 21     2  16   14 1_naturalistic
## 31     3  11    8 1_naturalistic
## 41     4  13   10 1_naturalistic
## 51     5  13   12 1_naturalistic
## 61     6   4   13 1_naturalistic
## 81     8   9   13 1_naturalistic
## 91     9   9    9 1_naturalistic
## 101   10  16   12 1_naturalistic
## 11    11  12   14 1_naturalistic
## 12    12  12    7 1_naturalistic
## 14     1  11   12  2_pentadecane
## 22     2  12    5  2_pentadecane
## 32     3   9   18  2_pentadecane
## 42     4  16   10  2_pentadecane
## 52     5   6   12  2_pentadecane
## 62     6  13   10  2_pentadecane
## 71     7   9   18  2_pentadecane
## 82     8  15   12  2_pentadecane
## 92     9   7   14  2_pentadecane
## 102   10  21    9  2_pentadecane
## 111   11  13    6  2_pentadecane
## 121   12   4   11  2_pentadecane
```

#### Turn the data into binary format

Each pupa in the 'stim' [stimulus] category counts as a success ('1') and each pupa in the 'con' [control] category counts as a failure ('0'). Stimulus and control are defined as follows for the three experimental conditions:

Substrate: Stimulus = leaf litter, Control = sand; Pentadecane: Stimulus = pentadecane, Control = paraffin oil; Naturalistic: Stimulus = conditioned sand (20 pupae), Control = unconditioned sand

```
data |> mutate(binary = map2(stim, con,
                                   ~ c(rep(1, .x),
                                       rep(0, .y)))) -> data_binary
data_binary <- unnest(data=data_binary, cols=binary)
class(data_binary$cage) <- 'factorial'
class(data_binary$experiment) <- 'factorial'
data_binary
```

```
## # A tibble: 751 x 5
##    cage          con  stim experiment  binary
##    <factoril> <int> <int> <factoril>   <dbl>
##  1 1              2    15 3_substrate      1
##  2 1              2    15 3_substrate      1
##  3 1              2    15 3_substrate      1
##  4 1              2    15 3_substrate      1
##  5 1              2    15 3_substrate      1
##  6 1              2    15 3_substrate      1
##  7 1              2    15 3_substrate      1
##  8 1              2    15 3_substrate      1
##  9 1              2    15 3_substrate      1
## 10 1              2    15 3_substrate      1
## # ... with 741 more rows
## # i Use `print(n = ...)` to see more rows
```

#### Prepare brms model

with 'cage' as group-level effect and 'experiment' as population-level effect. Adjust beta priors from flat to normal.

```
formula0 <- brms::brmsformula('binary ~ experiment + (1|cage)')
prior_b <- prior(normal(0,5), class = b)
formula0
```

```
## binary ~ experiment + (1 | cage)
```

```
prior_b
```

```
## b ~ normal(0, 5)
```

As the dependent variable is binary, samples will be drawn from a Bernoulli distribution with logit link.

```
## Compiling Stan program...
```

```
## Trying to compile a simple C file

## Running /Library/Frameworks/R.framework/Resources/bin/R CMD SHLIB foo.c
## clang -mmacosx-version-min=10.13 -I"/Library/Frameworks/R.framework/Resources/include" -DNDEBUG   -I
## In file included from <built-in>:1:
## In file included from /Library/Frameworks/R.framework/Versions/4.2/Resources/library/StanHeaders/incl
## In file included from /Library/Frameworks/R.framework/Versions/4.2/Resources/library/RcppEigen/inclu
## In file included from /Library/Frameworks/R.framework/Versions/4.2/Resources/library/RcppEigen/inclu
## /Library/Frameworks/R.framework/Versions/4.2/Resources/library/RcppEigen/include/Eigen/src/Core/util/
## namespace Eigen {
## ^
## /Library/Frameworks/R.framework/Versions/4.2/Resources/library/RcppEigen/include/Eigen/src/Core/util/
## namespace Eigen {
##                ^
##                ;
## In file included from <built-in>:1:
## In file included from /Library/Frameworks/R.framework/Versions/4.2/Resources/library/StanHeaders/incl
## In file included from /Library/Frameworks/R.framework/Versions/4.2/Resources/library/RcppEigen/inclu
## /Library/Frameworks/R.framework/Versions/4.2/Resources/library/RcppEigen/include/Eigen/Core:96:10: fa
## #include <complex>
##          ^~~~~~~~~
## 3 errors generated.
## make: *** [foo.o] Error 1

## Start sampling

##
## SAMPLING FOR MODEL '2fbc18f6c27e8d76d359b0d57eea1b46' NOW (CHAIN 1).
## Chain 1:
## Chain 1: Gradient evaluation took 9.5e-05 seconds
## Chain 1: 1000 transitions using 10 leapfrog steps per transition would take 0.95 seconds.
## Chain 1: Adjust your expectations accordingly!
## Chain 1:
## Chain 1:
## Chain 1: Iteration:    1 / 2000 [  0%]  (Warmup)
## Chain 1: Iteration:  200 / 2000 [ 10%]  (Warmup)
## Chain 1: Iteration:  400 / 2000 [ 20%]  (Warmup)
## Chain 1: Iteration:  600 / 2000 [ 30%]  (Warmup)
## Chain 1: Iteration:  800 / 2000 [ 40%]  (Warmup)
## Chain 1: Iteration: 1000 / 2000 [ 50%]  (Warmup)
## Chain 1: Iteration: 1001 / 2000 [ 50%]  (Sampling)
## Chain 1: Iteration: 1200 / 2000 [ 60%]  (Sampling)
## Chain 1: Iteration: 1400 / 2000 [ 70%]  (Sampling)
## Chain 1: Iteration: 1600 / 2000 [ 80%]  (Sampling)
## Chain 1: Iteration: 1800 / 2000 [ 90%]  (Sampling)
## Chain 1: Iteration: 2000 / 2000 [100%]  (Sampling)
## Chain 1:
## Chain 1:  Elapsed Time: 0.682764 seconds (Warm-up)
## Chain 1:                0.647232 seconds (Sampling)
## Chain 1:                1.33 seconds (Total)
## Chain 1:
##
## SAMPLING FOR MODEL '2fbc18f6c27e8d76d359b0d57eea1b46' NOW (CHAIN 2).
```

```
## Chain 2:
## Chain 2: Gradient evaluation took 4.9e-05 seconds
## Chain 2: 1000 transitions using 10 leapfrog steps per transition would take 0.49 seconds.
## Chain 2: Adjust your expectations accordingly!
## Chain 2:
## Chain 2:
## Chain 2: Iteration:    1 / 2000 [  0%]  (Warmup)
## Chain 2: Iteration:  200 / 2000 [ 10%]  (Warmup)
## Chain 2: Iteration:  400 / 2000 [ 20%]  (Warmup)
## Chain 2: Iteration:  600 / 2000 [ 30%]  (Warmup)
## Chain 2: Iteration:  800 / 2000 [ 40%]  (Warmup)
## Chain 2: Iteration: 1000 / 2000 [ 50%]  (Warmup)
## Chain 2: Iteration: 1001 / 2000 [ 50%]  (Sampling)
## Chain 2: Iteration: 1200 / 2000 [ 60%]  (Sampling)
## Chain 2: Iteration: 1400 / 2000 [ 70%]  (Sampling)
## Chain 2: Iteration: 1600 / 2000 [ 80%]  (Sampling)
## Chain 2: Iteration: 1800 / 2000 [ 90%]  (Sampling)
## Chain 2: Iteration: 2000 / 2000 [100%]  (Sampling)
## Chain 2:
## Chain 2:  Elapsed Time: 0.707563 seconds (Warm-up)
## Chain 2:                0.659292 seconds (Sampling)
## Chain 2:                1.36686 seconds (Total)
## Chain 2:
##
## SAMPLING FOR MODEL '2fbc18f6c27e8d76d359b0d57eea1b46' NOW (CHAIN 3).
## Chain 3:
## Chain 3: Gradient evaluation took 5.4e-05 seconds
## Chain 3: 1000 transitions using 10 leapfrog steps per transition would take 0.54 seconds.
## Chain 3: Adjust your expectations accordingly!
## Chain 3:
## Chain 3:
## Chain 3: Iteration:    1 / 2000 [  0%]  (Warmup)
## Chain 3: Iteration:  200 / 2000 [ 10%]  (Warmup)
## Chain 3: Iteration:  400 / 2000 [ 20%]  (Warmup)
## Chain 3: Iteration:  600 / 2000 [ 30%]  (Warmup)
## Chain 3: Iteration:  800 / 2000 [ 40%]  (Warmup)
## Chain 3: Iteration: 1000 / 2000 [ 50%]  (Warmup)
## Chain 3: Iteration: 1001 / 2000 [ 50%]  (Sampling)
## Chain 3: Iteration: 1200 / 2000 [ 60%]  (Sampling)
## Chain 3: Iteration: 1400 / 2000 [ 70%]  (Sampling)
## Chain 3: Iteration: 1600 / 2000 [ 80%]  (Sampling)
## Chain 3: Iteration: 1800 / 2000 [ 90%]  (Sampling)
## Chain 3: Iteration: 2000 / 2000 [100%]  (Sampling)
## Chain 3:
## Chain 3:  Elapsed Time: 0.881934 seconds (Warm-up)
## Chain 3:                0.678404 seconds (Sampling)
## Chain 3:                1.56034 seconds (Total)
## Chain 3:
##
## SAMPLING FOR MODEL '2fbc18f6c27e8d76d359b0d57eea1b46' NOW (CHAIN 4).
## Chain 4:
## Chain 4: Gradient evaluation took 7e-05 seconds
## Chain 4: 1000 transitions using 10 leapfrog steps per transition would take 0.7 seconds.
## Chain 4: Adjust your expectations accordingly!
```

```
## Chain 4:
## Chain 4:
## Chain 4: Iteration:    1 / 2000 [  0%]  (Warmup)
## Chain 4: Iteration:  200 / 2000 [ 10%]  (Warmup)
## Chain 4: Iteration:  400 / 2000 [ 20%]  (Warmup)
## Chain 4: Iteration:  600 / 2000 [ 30%]  (Warmup)
## Chain 4: Iteration:  800 / 2000 [ 40%]  (Warmup)
## Chain 4: Iteration: 1000 / 2000 [ 50%]  (Warmup)
## Chain 4: Iteration: 1001 / 2000 [ 50%]  (Sampling)
## Chain 4: Iteration: 1200 / 2000 [ 60%]  (Sampling)
## Chain 4: Iteration: 1400 / 2000 [ 70%]  (Sampling)
## Chain 4: Iteration: 1600 / 2000 [ 80%]  (Sampling)
## Chain 4: Iteration: 1800 / 2000 [ 90%]  (Sampling)
## Chain 4: Iteration: 2000 / 2000 [100%]  (Sampling)
## Chain 4:
## Chain 4:  Elapsed Time: 0.761918 seconds (Warm-up)
## Chain 4:                0.644089 seconds (Sampling)
## Chain 4:                1.40601 seconds (Total)
## Chain 4:
```

```
##  Family: bernoulli
##   Links: mu = logit
## Formula: binary ~ experiment + (1 | cage)
##    Data: data_binary (Number of observations: 751)
##   Draws: 4 chains, each with iter = 2000; warmup = 1000; thin = 1;
##          total post-warmup draws = 4000
##
## Group-Level Effects:
## ~cage (Number of levels: 12)
##               Estimate Est.Error l-95% CI u-95% CI Rhat Bulk_ESS Tail_ESS
## sd(Intercept)     0.20      0.13     0.01     0.51 1.00     1218     1320
##
## Population-Level Effects:
##                        Estimate Est.Error l-95% CI u-95% CI Rhat Bulk_ESS
## Intercept                  0.05      0.15    -0.23     0.34 1.00     2888
## experiment2_pentadecane   -0.05      0.18    -0.41     0.30 1.00     4577
## experiment3_substrate      2.23      0.27     1.70     2.77 1.00     3634
##                        Tail_ESS
## Intercept                  1809
## experiment2_pentadecane    3089
## experiment3_substrate      2722
##
## Draws were sampled using sampling(NUTS). For each parameter, Bulk_ESS
## and Tail_ESS are effective sample size measures, and Rhat is the potential
## scale reduction factor on split chains (at convergence, Rhat = 1).
```

#### Evaluate model

Rhat and Bulk_ESS values suggest that the model converged well.

To confirm, we plot the model to visually inspect the chains.

```r
plot(data_m0)
```

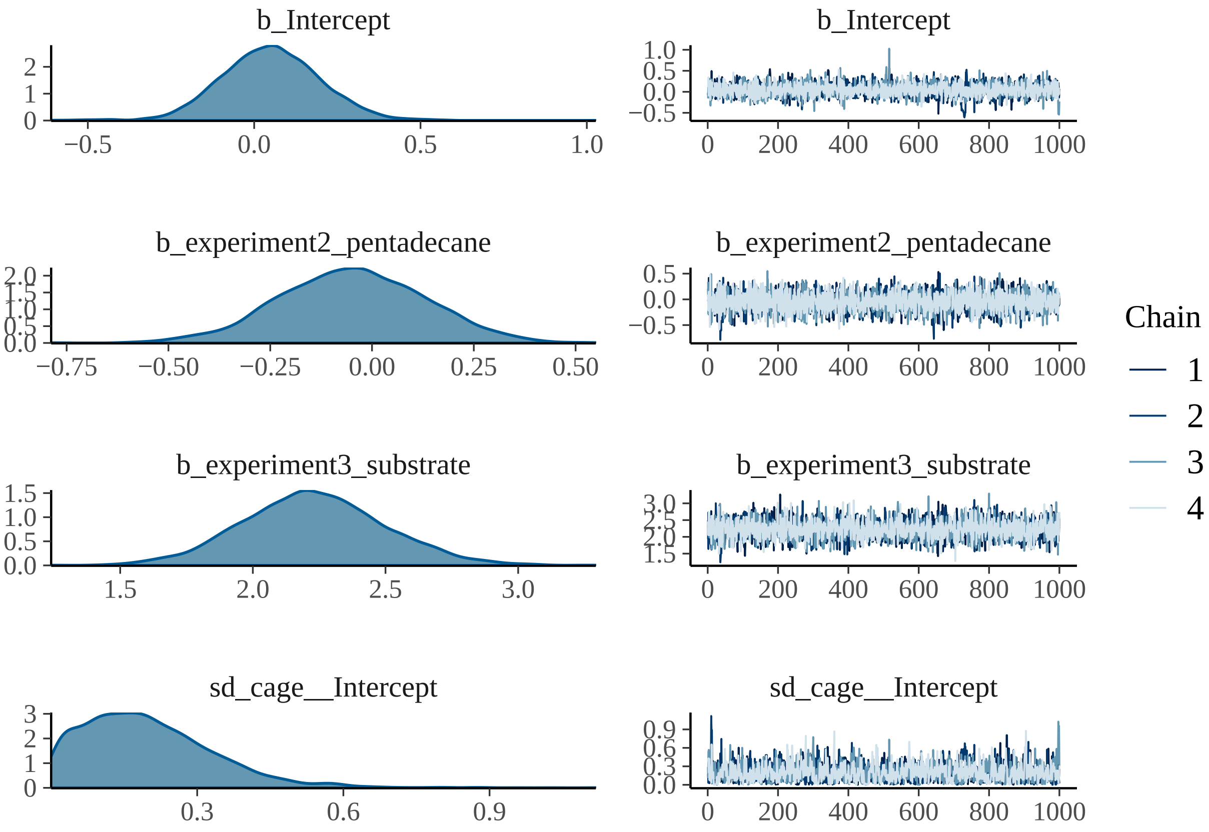


Chains look good, the model has converged well.

#### Model results

We can now look at conditional effects predicted by the model.

```r
p <- conditional_effects(data_m0)
plot(p)
```

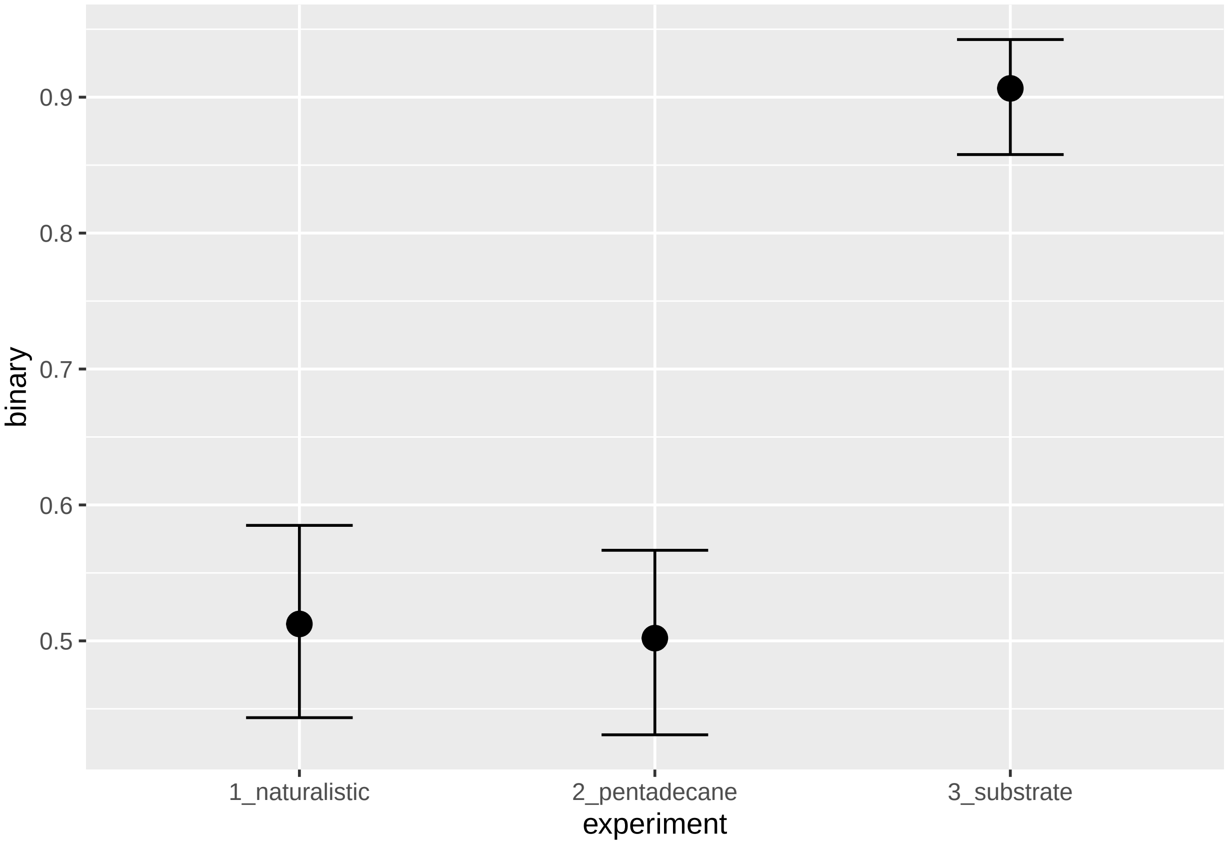


```r
p[[1]]
```

```
##       experiment    binary cage cond__      effect1__ estimate__       se__
## 1 1_naturalistic 0.6218375   NA      1 1_naturalistic  0.5124503 0.03581315
## 2  2_pentadecane 0.6218375   NA      1  2_pentadecane  0.5020129 0.03385076
## 3    3_substrate 0.6218375   NA      1    3_substrate  0.9064589 0.02030241
##     lower__   upper__
## 1 0.4434994 0.5849641
## 2 0.4308909 0.5666415
## 3 0.8577875 0.9424012
```

The model supports a preference only in the 'substrate' condition, with an estimated preference of 0.9067 for '1', i.e. leaf litter. No preference is supported for the 'pentadecane' and 'naturalistic' experiments. Thus, the conditional effects support the results from behavioural experiments and mirror the preference index analysis.

#### Hypothesis tests

Just to make sure, we can check the outcomes for each individual population- level effect:

```r
hyp1 <- hypothesis(data_m0, 'Intercept > 0')
plot(hyp1)
```

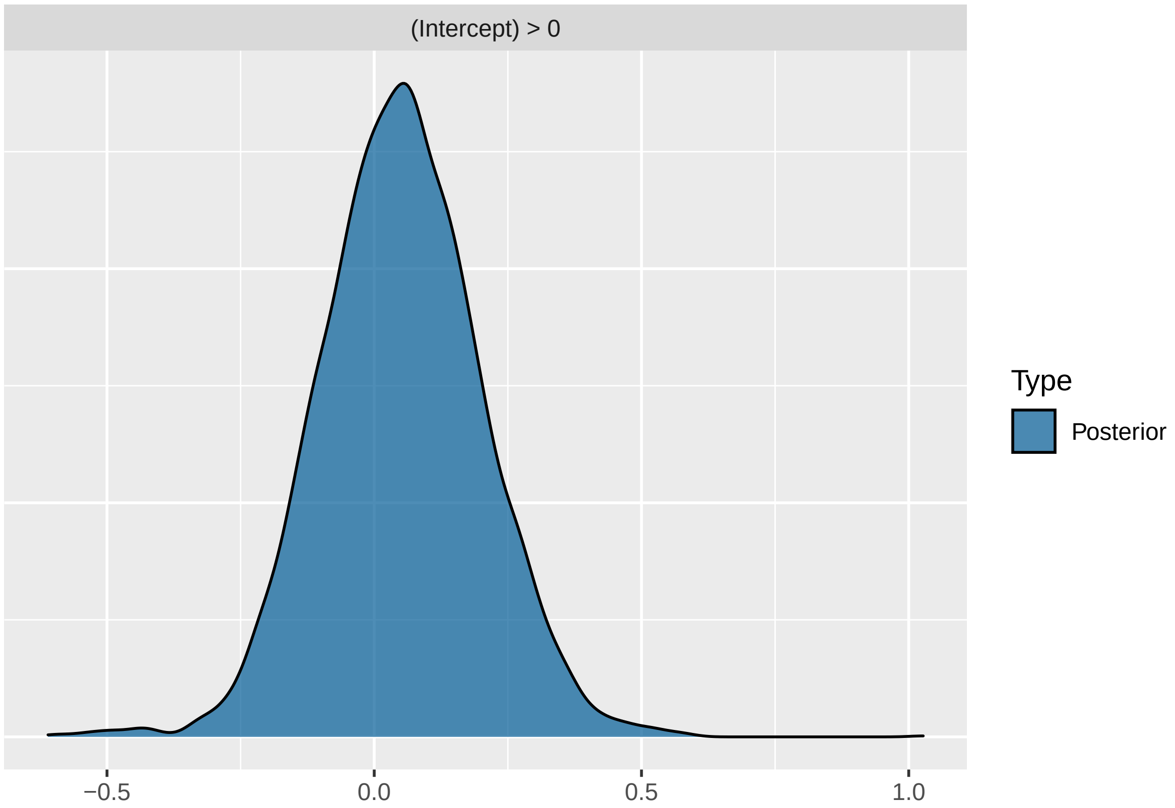


```
hyp1
```

```
## Hypothesis Tests for class b:
##        Hypothesis Estimate Est.Error CI.Lower CI.Upper Evid.Ratio Post.Prob
## 1 (Intercept) > 0     0.05      0.15    -0.19     0.29       1.71      0.63
##   Star
## 1
## ---
## 'CI': 90%-CI for one-sided and 95%-CI for two-sided hypotheses.
## '*': For one-sided hypotheses, the posterior probability exceeds 95%;
## for two-sided hypotheses, the value tested against lies outside the 95%-CI.
## Posterior probabilities of point hypotheses assume equal prior probabilities.
```

Intercept is a dummy variable for the 'naturalistic' condition. The distribution has an estimated median of 0.06 and does not support a preference for either side.

```
hyp2 <- hypothesis(data_m0, 'experiment2_pentadecane > 0')
plot(hyp2)
```

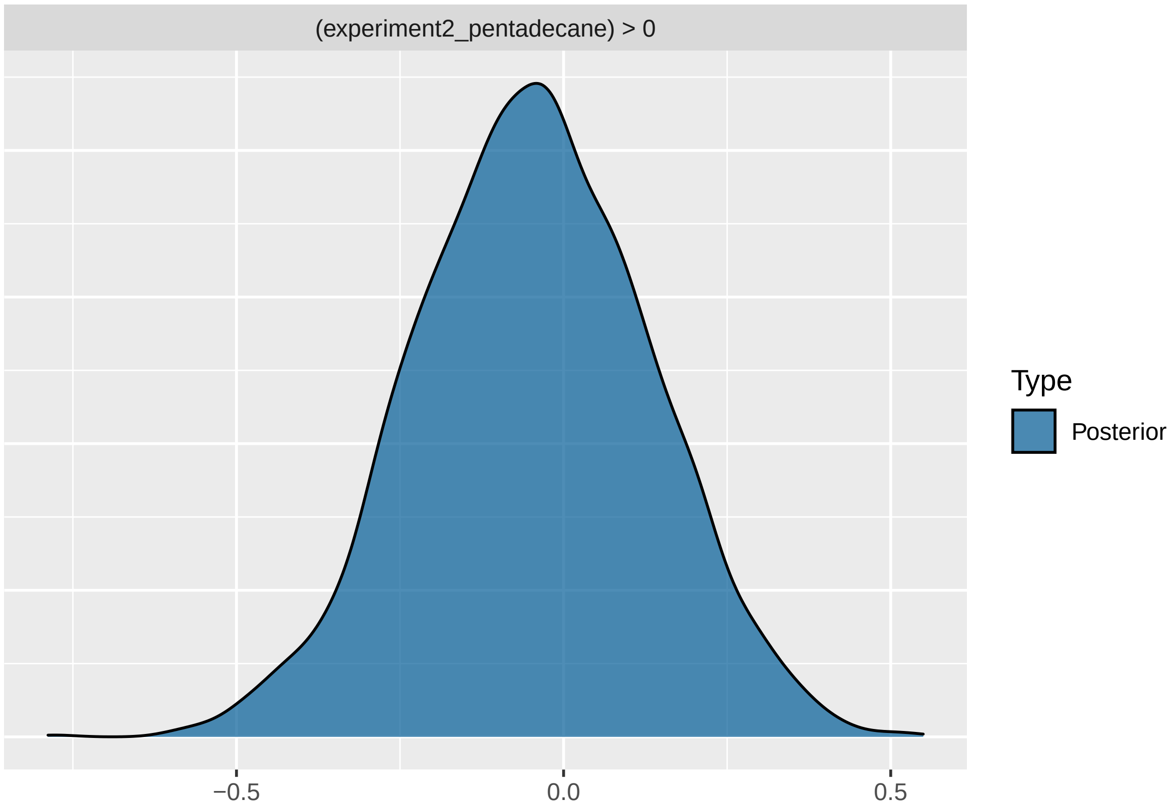


```
hyp2
```

```
## Hypothesis Tests for class b:
##                 Hypothesis Estimate Est.Error CI.Lower CI.Upper Evid.Ratio
## 1 (experiment2_pent... > 0    -0.05      0.18    -0.34     0.24       0.64
##   Post.Prob Star
## 1      0.39
## ---
## 'CI': 90%-CI for one-sided and 95%-CI for two-sided hypotheses.
## '*': For one-sided hypotheses, the posterior probability exceeds 95%;
## for two-sided hypotheses, the value tested against lies outside the 95%-CI.
## Posterior probabilities of point hypotheses assume equal prior probabilities.
```

The distribution for the 'pentadecane' condition has an estimate of -0.06 and does not support a preference for either side.

```
hyp3 <- hypothesis(data_m0, 'experiment3_substrate > 0')
plot(hyp3)
```

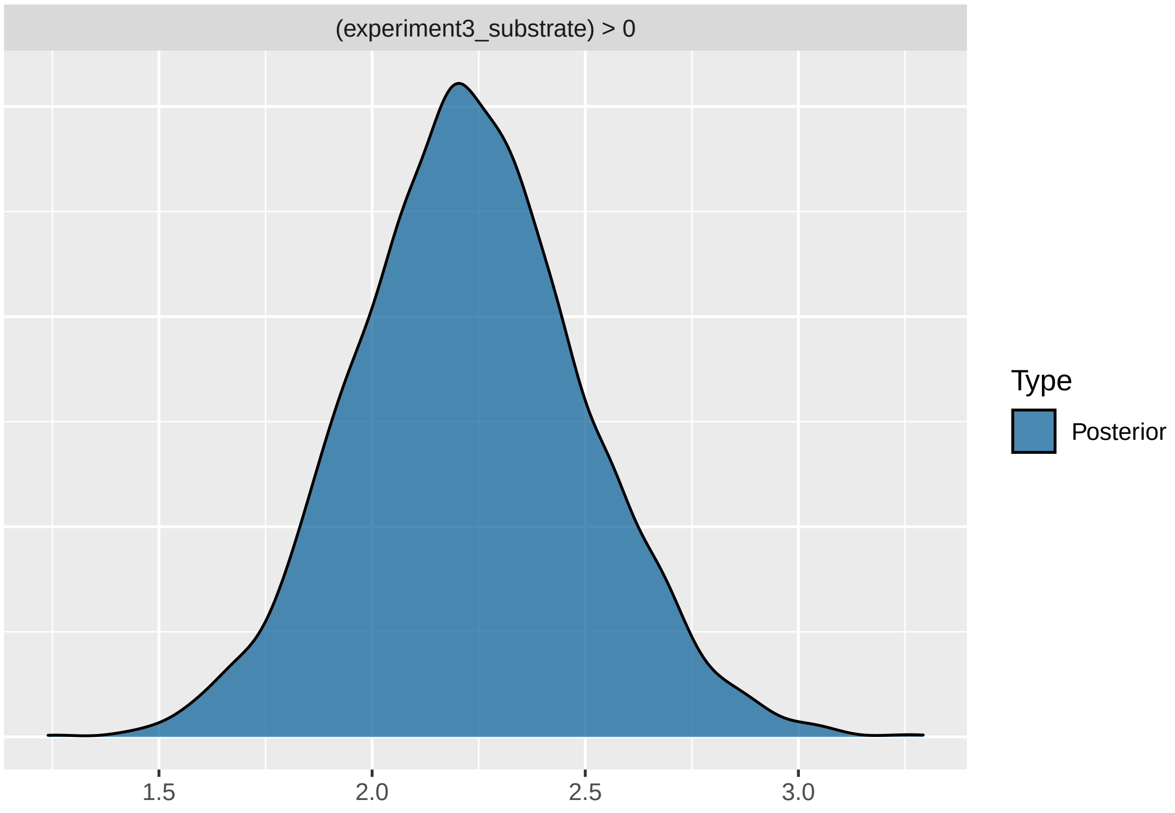


```
hyp3
```

```
## Hypothesis Tests for class b:
##                Hypothesis Estimate Est.Error CI.Lower CI.Upper Evid.Ratio
## 1 (experiment3_subs... > 0     2.23      0.27      1.8     2.68        Inf
##   Post.Prob Star
## 1         1    *
## ---
## 'CI': 90%-CI for one-sided and 95%-CI for two-sided hypotheses.
## '*': For one-sided hypotheses, the posterior probability exceeds 95%;
## for two-sided hypotheses, the value tested against lies outside the 95%-CI.
## Posterior probabilities of point hypotheses assume equal prior probabilities.
```

The distribution for the 'substrate' condition clusters around an estimate of 2.22 with an infinite evidence ratio and a posterior probability of 1, supporting a strong preference for choosing '1', i.e. leaf litter.
